# Characterization of Lactobacillus rhamnosus-Derived Extracellular Vesicles and Their Cellular Cytoprotective Effects Against Trimethylamine-Induced Oxidative Stress in Caco-2 Cells

**DOI:** 10.64898/2026.11.04.764202

**Authors:** Le-Ming Wang, Then Chee Kin, Wei-Chia Hung, Yeu-Ching Shi

**Affiliations:** Department of Obstetrics and Gynecology, Wan Fang Hospital, Taipei Medical University, Taipei, Taiwan; The College of Medicine of Taipei Medical University, Taipei, Taiwan; Bio X Biotech Co., Ltd, Taipei, Taiwan

**Keywords:** *Lactobacillus rhamnosus*, Extracellular vesicles, Trimethylamine (TMA), SIRT1 pathway, Zonula Occludens-1 (ZO-1)

## Abstract

Dietary trimethylamine (TMA) underwent gut microbiota-mediated catabolism and subsequent hepatic oxidation to generate trimethylamine N-oxide (TMAO), a key metabolic factor linked to vascular and colorectal diseases. While the systemic effects of circulating TMAO were heavily investigated, the direct localized cellular interactions of its precursor, TMA, at the intestinal epithelial level required further elucidation. This study characterized extracellular vesicles isolated from *Lactobacillus rhamnosus* (Lr-EVs) and evaluated their cytoprotective potential in an *in vitro* human intestinal Caco-2 cell model under TMA-induced stress. Exposure to a high concentration of TMA (20 mM) significantly reduced Caco-2 viability to 78.8%, triggered intracellular reactive oxygen species (ROS) accumulation, and compromised tight junction integrity as observed via the disruption of Zonula Occludens-1 (ZO-1) alignment. Co-treatment with Lr-EVs (100-300 μg/mL) counteracted this cellular toxicity, enhancing cell survival up to 172%. Quantitative PCR analysis revealed that Lr-EVs suppressed the transcriptional upregulation of the pro-inflammatory gene IL-1β, restored the expression of the metabolic regulator SIRT1, and stabilized transcript levels of key mitochondrial-encoded electron transport chain genes (ND6, CYTB, CO1, and ATP6). Additionally, Lr-EVs exhibited intrinsic radical scavenging activities in cell-free ABTS and DPPH assays. These preliminary *in vitro* findings demonstrated that Lr-EVs mitigated TMA-induced cellular injury, oxidative stress, and junctional disruption at the transcriptomic level, supporting further investigation of probiotic-derived EVs as prospective cell-free functional ingredients.

**Highlights:**

- Successful isolation of functional extracellular vesicles from *Lactobacillus rhamnosus* was achieved.
- Lr-EVs rescued Caco-2 cells from high-dose trimethylamine-induced localized cytotoxicity.
- Lr-EVs suppressed oxidative stress, inflammation, and mitochondrial transcript degradation.
- The cytoprotective effects were correlated with the transcriptional restoration of the SIRT1 pathway and ZO-1 alignment preservation.

## Introduction

The demographic shift toward a globally aging population has dramatically escalated the incidence of age-related metabolic and chronic inflammatory disorders, including cardiovascular disease (CVD) and chronic kidney disease (CKD) (1). Emerging evidence highlights the gut microbiota and its small-molecule metabolome as central arbiters of host metabolic homeostasis and disease progression. Within this microbial metabolic network, the trimethylamine (TMA)-trimethylamine N-oxide (TMAO) pathway represents a critical pathogenic axis. Intestinal commensals catabolize dietary quaternary amines-such as choline, L-carnitine, and betaine-into volatile TMA, which is subsequently absorbed across the epithelium and oxidized by hepatic flavin-containing monooxygenases (FMOs) to form systemic TMAO (2, 3).

While a substantial body of literature has elucidated the systemic pro-atherogenic and pro-thrombotic mechanisms of circulating TMAO, the immediate localized effects of its precursor, TMA, on the intestinal epithelial interface remain enigmatic. In elderly populations, progressive age-associated gut dysbiosis-characterized by the expansion of distinct phyla such as *Pseudomonadota* and genera like *Prevotella*-accelerates luminal TMA accumulation. Exposure to elevated concentrations of TMA induces pronounced localized epithelial stress, characterized by the excessive generation of reactive oxygen species (ROS), mitochondrial dysfunction, and structural impairment of the intestinal mucosal barrier (4–6). This compromised epithelial barrier creates a permissive environment for accelerated solute translocation, intensifying the chronic low-grade inflammation linked to colorectal carcinogenesis and driving systemic metabolic toxemia (7). Epidemiological cohorts indicate that the average plasma TMAO level is approximately 2.8 mM in younger adults (aged 18-44), increases to 4.4 mM in middle-aged individuals (aged 45-64), and reaches 9.8 mM in healthy elderly individuals over 65 years old (8). Furthermore, postmenopausal women aged over 55 present a higher risk of developing colorectal cancer (CRC) compared to premenopausal women under 40, suggesting that hormonal transitions combined with age-related gut dysbiosis accelerate metabolic toxicity (9). Consequently, intercepting the toxic cascade of TMA directly at the mucosal interface presents a promising therapeutic approach for mitigating downstream systemic pathologies.

Concurrently, cell-free postbiotic modalities have attracted scientific interest as alternatives to traditional live probiotics, which face challenges regarding viable gastrointestinal transit and complex colonization kinetics in competitive host niches (10, 11). Studies have demonstrated that *Lactobacillus salivarius* has the potential to prevent *Salmonella* infection, and the supplementation of *Bifidobacterium* and cellulose can inhibit the overgrowth of *Clostridium difficile* (12, 13). In addition to competitive exclusion mechanisms, recent breakthroughs in microbial endocrinology have shown that gut microbes communicate with host cells by modulating host microRNAs (miRNAs), thereby reducing inflammation and endothelial damage (14, 15). *Lactobacillus plantarum* ZDY04 has been shown to enhance cecal microbiota composition, thereby reducing cecal TMA and serum TMAO concentrations in murine models (16, 17). Probiotic-derived extracellular vesicles (EVs)-lipid-bilayer nano-sized structures carrying bioactive proteins, lipids, and nucleic acids-serve as functional vectors for trans-kingdom communication (18). These microbial EVs function as delivery vehicles for bioactive proteins, lipopolysaccharides, and small non-coding RNAs. Recent evidence confirms that host fecal EVs rich in specific miRNAs can selectively promote or suppress the growth of specific intestinal microbes like *Fusobacterium nucleatum* and *Escherichia coli*, demonstrating a regulatory axis that maintains the gut ecosystem (19). Although specific probiotic strains have been shown to modulate the gut microbiota composition and lower systemic TMAO levels *in vivo*, the precise capacity of probiotic EVs to interface directly with host enterocytes and preserve mucosal homeostasis against metabolic insults remains unexplored.

To investigate these interaction dynamics, this *in vitro* study aimed to isolate, biophysically characterize, and evaluate the cytoprotective efficacy of EVs derived from lactic acid bacteria (LAB) strains. Using a standard Caco-2 cell culture system, we demonstrate that *Lactobacillus rhamnosus*-derived EVs (Lr-EVs) possess intrinsic antioxidant activities and protect cellular viability, transcriptomic mitochondrial stability, and tight junction arrangement against high-dose TMA challenges, potentially correlating with the upregulation of the SIRT1 pathway.

## Materials and Methods

### 2.1. Lactic acid bacteria culture and extracellular vesicle isolation

Lyophilized stocks of lactic acid bacteria (LAB) strains (provided by Bio X Biotech Co., Ltd., Taipei, Taiwan) were rehydrated and inoculated into de Man, Rogosa, and Sharpe (MRS) broth (Difco Laboratories, Detroit, MI, USA), followed by incubation at 37 °C for 12 h. Cultured bacteria were subsequently subcultured (1:100, v/v) into fresh MRS broth and monitored spectrophotometrically until reaching the mid-logarithmic growth phase (optical density at 600 nm, OD600 = 0.7). The cultures were maintained for an additional 24 h at 37 °C to optimize extracellular vesicle (EV) secretion. To isolate the microbial vesicles, the conditioned media were subjected to sequential differential centrifugation at 3,000 ×g for 15 min at 4 °C and 10,000 ×g for 30 min at 4 °C to remove intact bacterial cells and cellular debris. The resulting supernatant was passed through a 0.22 μm polyethersulfone membrane filter (Millipore, Bedford, MA, USA) and subjected to high-speed ultracentrifugation at 100,000 ×g for 2 h at 4 °C to sediment the nano-sized vesicles. The final EV pellets were resuspended in sterile Dulbecco’s phosphate-buffered saline (DPBS; Gibco, Grand Island, NY, USA) and stored at −80 °C until downstream usage.

### 2.2. Morphological and biophysical characterization of LAB-EVs

For ultrastructural examination, a 10 μL aliquot of the suspended vesicle sample was applied onto carbon-coated 400-mesh copper grids, negatively stained with 2% (w/v) uranyl acetate, and visualized via transmission electron microscopy (TEM; JEM-2100, JEOL, Tokyo, Japan). Size distribution profiles and absolute particle concentrations were precisely determined through nanoparticle tracking analysis (NTA) using a NanoSight NS300 instrument (Malvern Panalytical, Malvern, UK) after appropriate dilution in particle-free DPBS.

### 2.3. Quantification of EV biomolecular content

Total DNA associated with the purified LAB-EVs was quantified using a NanoDrop 2000 microvolume spectrophotometer (Thermo Fisher Scientific, Wilmington, DE, USA) via absorbance ratios (A260/A280). Total vesicular protein concentration was determined utilizing the Bradford colorimetric assay, with a standard calibration curve constructed from sequential dilutions of bovine serum albumin (BSA; Sigma-Aldrich, St. Louis, MO, USA). Aliquots of various concentrations of BSA standards or LAB-EV samples (20 μL) were transferred to a 96-well plate, followed by the addition of 200 μL of Bradford reagent (Bio-Rad Laboratories, Hercules, CA, USA). After incubation at room temperature (25 °C) for 10 min, the optical density was recorded at 595 nm using a microplate reader (Multiskan GO, Thermo Fisher Scientific, Vantaa, Finland). The protein concentration was calculated based on the resulting linear standard curve equation.

### 2.4. Cell culture and viability analysis

Human colorectal adenocarcinoma (Caco-2) cells were obtained from the American Type Culture Collection (ATCC, Manassas, VA, USA) and cultured in Dulbecco’s modified Eagle’s medium (DMEM; Gibco) supplemented with 10% (v/v) fetal bovine serum (FBS; Gibco) and 1% (v/v) penicillin-streptomycin in a humidified incubator containing 5% CO₂ at 37 °C. To evaluate cyto-protection, cells were seeded in 96-well plates at a density of (1×10⁴) cells/well. Upon achieving confluence, monolayers were co-treated with graded concentrations of trimethylamine (TMA; 5-20 mM; Sigma-Aldrich) and purified LAB-EVs (100-300 μg/mL) for 24 h. Cell metabolic activity and relative viability were quantified using the Cell Counting Kit-8 (CCK-8; Dojindo Molecular Technologies, Kumamoto, Japan); absorbance was measured at 450 nm using a microplate reader following a 1 h incubation at 37 °C.

### 2.5. RNA extraction, cDNA synthesis, and quantitative real-time PCR (qPCR)

Total RNA was isolated from treated Caco-2 monolayers using the Total RNA Spin Kit (Geneaid Biotech, Taipei, Taiwan) following the manufacturer’s protocol. RNA purity and concentration were verified spectrophotometrically. Complementary DNA (cDNA) synthesis was executed using the ExcelRT Reverse Transcription Kit (GeneDireX, Taichung, Taiwan) under optimized thermal profiles according to the instructions. Quantitative real-time PCR (qPCR) was conducted using the Maxima SYBR Green/ROX qPCR Master Mix (Thermo Fisher Scientific) on an Applied Biosystems StepOnePlus Real-Time PCR System (Applied Biosystems, Foster City, CA, USA). Target gene transcription levels, including interleukin-1beta (IL-1β), Sirtuin 1 (SIRT1), and mitochondrial-encoded electron transport chain components (ND6, CYTB, CO1, and ATP6), were normalized to the endogenous reference gene glyceraldehyde-3-phosphate dehydrogenase (GAPDH) using the comparative calculation method.

### 2.6. Cell-free and intracellular antioxidant capacity assays

The cell-free radical-scavenging performance of LAB-EVs was validated using 2,2-diphenyl-1-picrylhydrazyl (DPPH) and 2,2’-azino-bis(3-ethylbenzothiazoline-6-sulfonic acid) radical cation assays as previously reported. To monitor intracellular reactive oxygen species (ROS) dynamics, Caco-2 cells subjected to TMA and EV treatments were labeled with 10 μM of the fluorogenic probe 2′,7′-dichlorodihydrofluorescein diacetate (DCFH-DA; Sigma-Aldrich) for 30 min at 37 °C. Relative intracellular fluorescence intensity was captured and quantified via an inverted fluorescence microscope (Olympus IX73, Tokyo, Japan) equipped with ImageJ software (version 1.54, National Institutes of Health, Bethesda, MD, USA).

### 2.7. Immunofluorescence staining for barrier assessment

To assess tight junction structural layout, Caco-2 monolayers cultured on glass coverslips were fixed with 4% (w/v) paraformaldehyde for 15 min, permeabilized with 0.1% (v/v) Triton X-100 for 10 min, and blocked with 3% (w/v) BSA for 1 h at room temperature. Cells were incubated with a primary rabbit anti-ZO-1 polyclonal antibody (1:200; Invitrogen, Carlsbad, CA, USA) overnight at 4 °C, followed by detection with an Alexa Fluor 488-conjugated goat anti-rabbit IgG secondary antibody (1:500; Invitrogen) for 1 h in the dark. The architectural arrangement and spatial distribution of Zonula Occludens-1 (ZO-1) along the cell borders were visualized via a high-resolution laser scanning confocal microscope (LSM 800, Carl Zeiss, Oberkochen, Germany).

### 2.8. Statistical analysis

All data are expressed as the mean ± standard deviation (SD) from three independent experiments. Statistical analysis was performed using the Statistical Analysis System (SAS) software package (version 9.4, SAS Institute Inc., Cary, NC, USA). Statistical significance among group means was evaluated by one-way analysis of variance (ANOVA) followed by Duncan’s multiple range test, or via a two-tailed Student’s t-test for single pairwise comparisons. P-values less than 0.05 (P < 0.05) were considered statistically significant.

## Results

### Biophysical Profiling and Selection of Probiotic LAB-EVs

Extracellular vesicles were successfully isolated from the spent media of *Lactobacillus rhamnosus* and *Lactobacillus lactis* subsp. *cremoris*. NTA characterization revealed distinct biophysical phenotypes: *L. rhamnosus*-derived EVs (Lr-EVs) displayed a mean hydrodynamic diameter of 94 nm, whereas *L. lactis*-derived EVs (Ll-EVs) were larger, averaging 123 nm. Crucially, *L. rhamnosus* demonstrated superior vesicle production kinetics, yielding a significantly higher particle concentration (1.13×10⁹ particles/mL; total protein: 4.70 mg/mL) relative to *L. lactis* (1.02×10⁸ particles/mL; total protein: 4.99 mg/mL). Given this robust secretory efficiency and high yield, Lr-EVs were selected as the primary candidate for all subsequent functional and mechanistic investigations (Fig 1).

**Figure 1.**
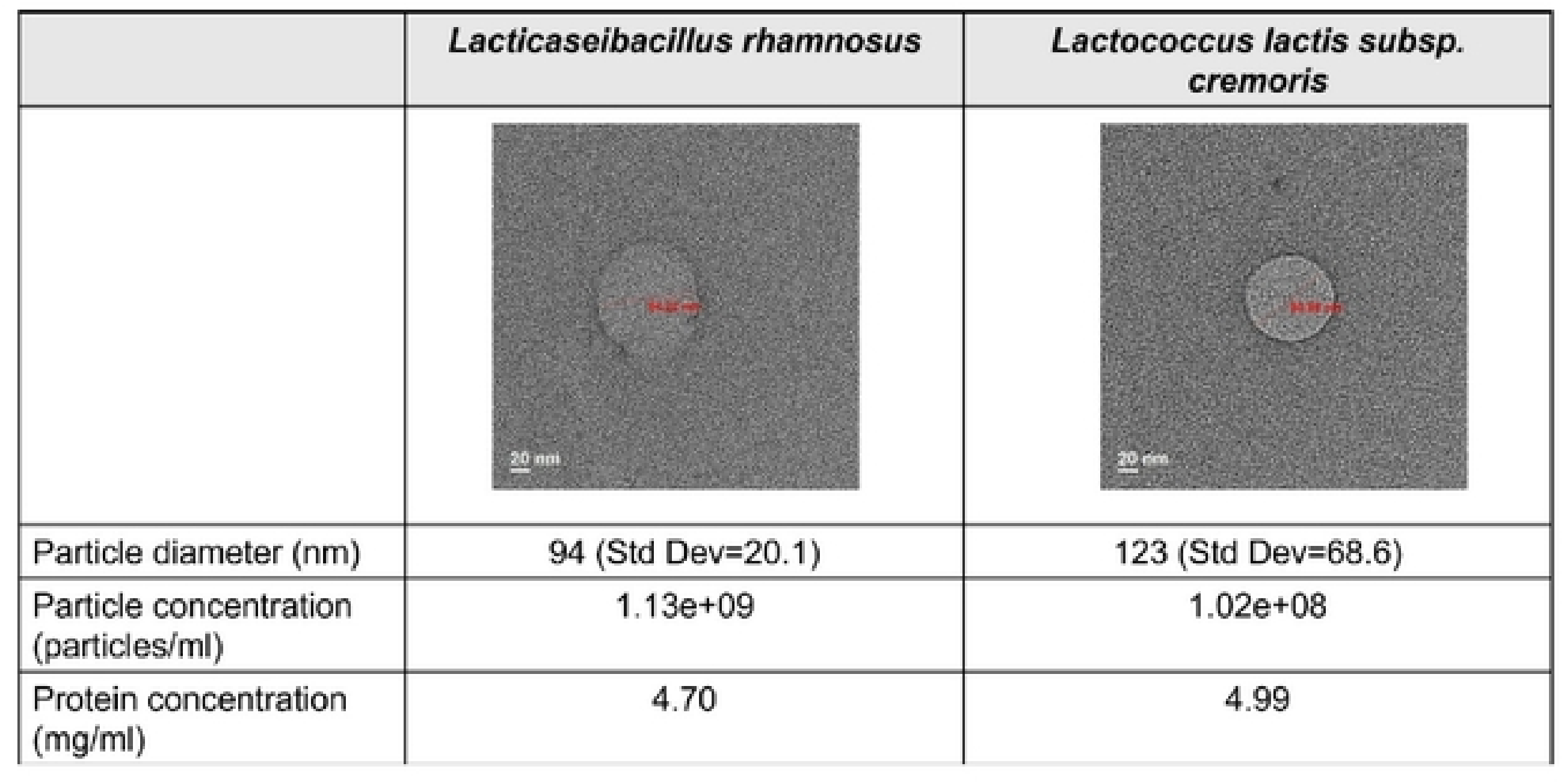
Biophysical characterization and isolation yield of lactic acid bacteria-derived extracellular vesicles (LAB-EVs).

### Lr-EVs Maintain Caco-2 Cell Viability Under TMA Challenge

To model localized metabolite-induced stress, Caco-2 cells were exposed to varying concentrations of TMA. Challenge with 20 mM TMA for 24 h induced a significant cytotoxic response, reducing cell viability to 78.8% of control levels. Remarkably, co-administration of Lr-EVs (100 and 300 μg/mL) not only completely neutralized TMA-induced lethality but profoundly boosted overall cell survival and proliferative capacity up to 172%. These baseline data suggest that Lr-EVs contain components that support cell survival and metabolic activity under high-dose chemical challenges *in vitro* (Fig 2).

**Figure 2.**
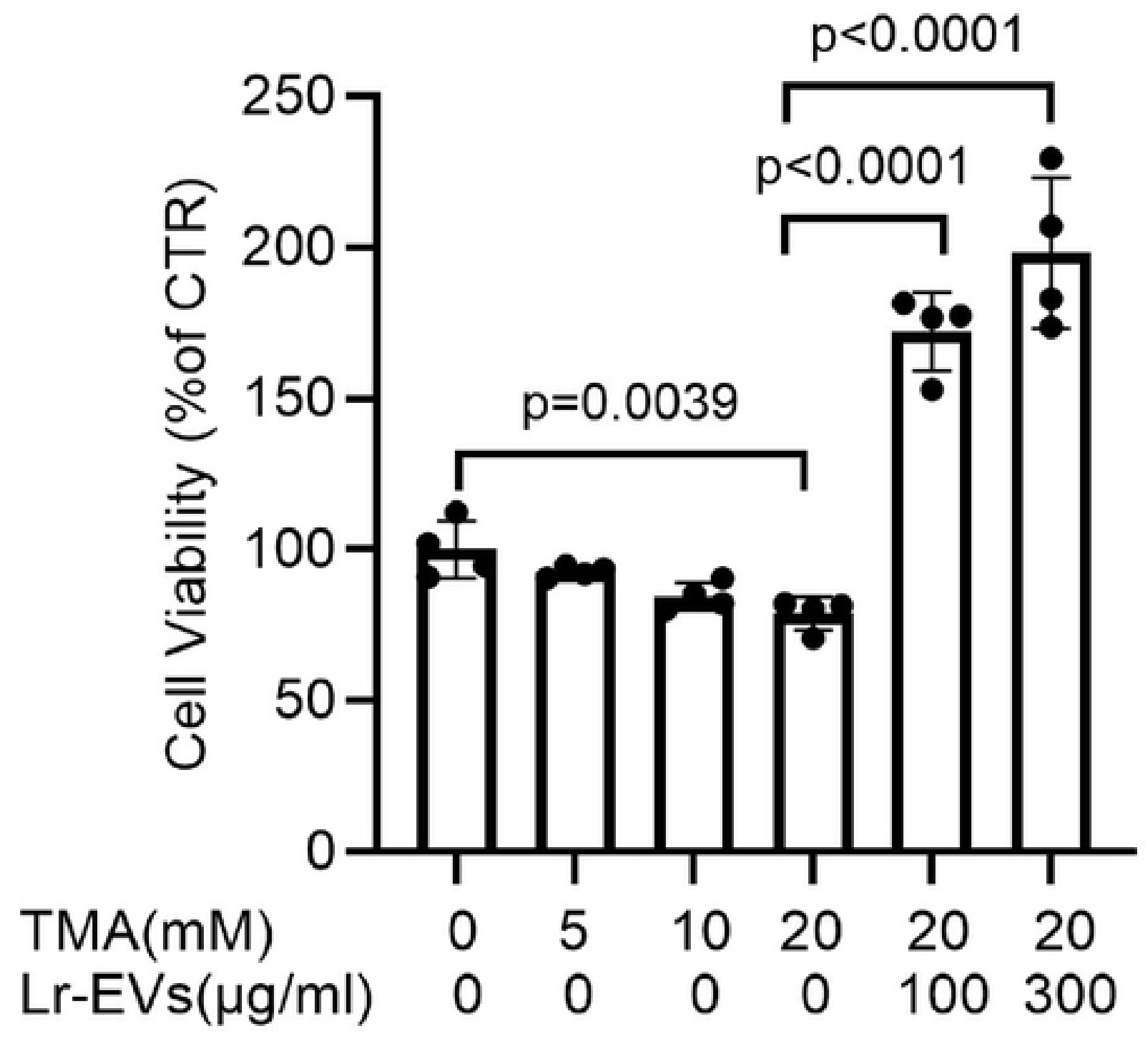
Protective efficacy of Lr-EVs against trimethylamine (TMA)-induced cytotoxicity in Caco-2 cells. Untreated cells served as the negative control. Values represent mean ± SD (n = 3). * p <0.05, **p<0.01,***p< 0.001 vs. untreated control group; ### p < 0.001 vs. TMA-treated group.

### Lr-EVs Co-treatment Modulates IL-1β and SIRT1 Transcript Levels

To assess the molecular changes associated with Lr-EV protection, qPCR was performed. TMA challenge triggered an inflammatory transcriptional response, marked by an upregulation of the pro-inflammatory cytokine IL-1β. Co-treatment with Lr-EVs reduced IL-1β mRNA accumulation in a dose-dependent manner. Lr-EV co-treatment successfully blunted this response, driving a dose-dependent downregulation of IL-1β transcript levels (Fig 3). Concurrently, transcript levels of Sirtuin 1 (SIRT1), a key metabolic regulator, were significantly downregulated following TMA exposure. Notably, the addition of Lr-EVs rescued and upregulated SIRT1 expression, suggesting that the cytoprotective effects of these vesicles may be linked to the transcriptional regulation of the SIRT1 pathway (Fig 3).

**Fig 3.**
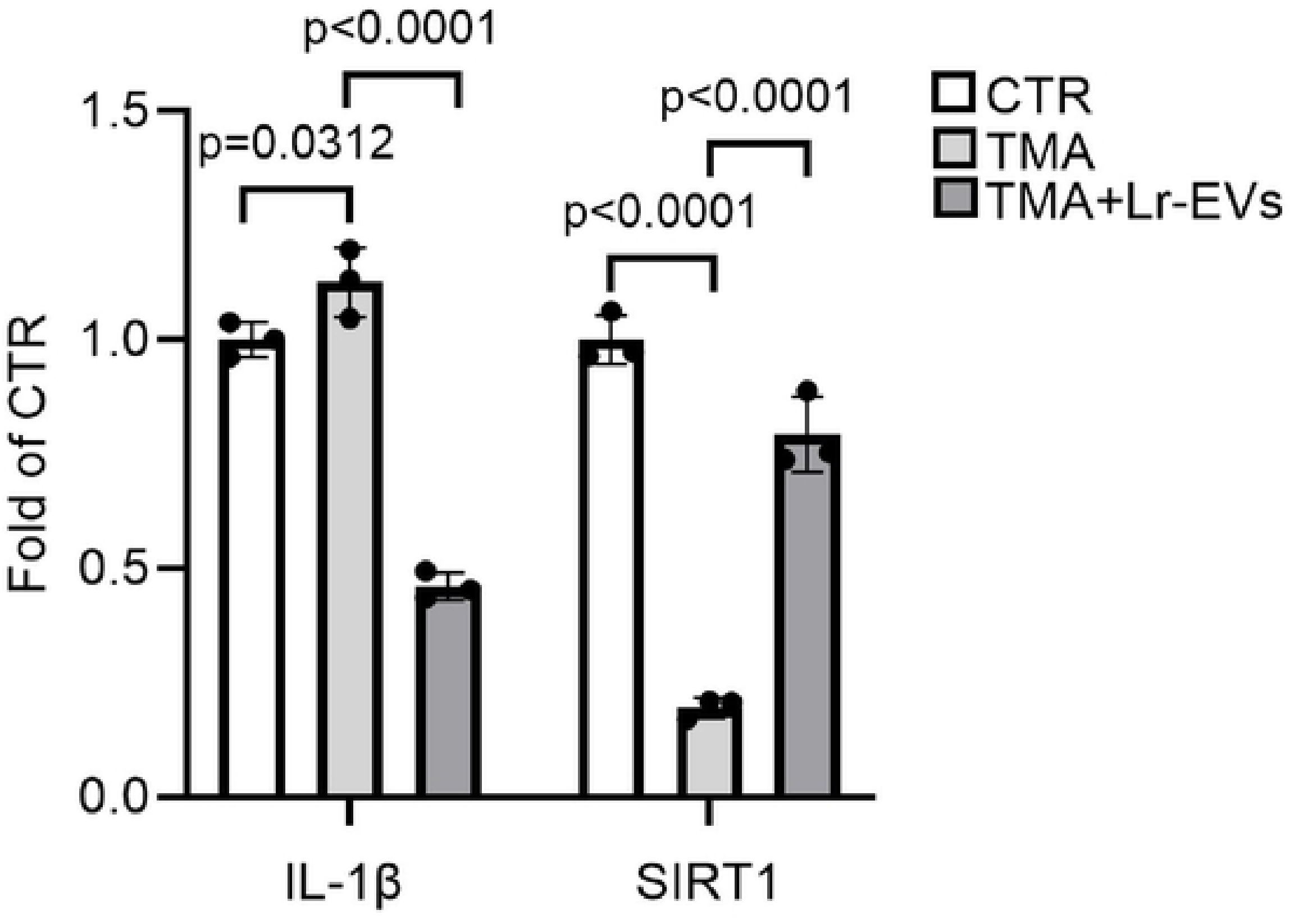
Lr-EVs suppress pro-inflammatory gene transcription and restore SIRT1 expression under TMA stress. mRNA expression levels of *SIRT1* determined by RT-qPCR analysis. Bar charts with individual data points (*n* = 3) represent mean ± SD. Exact P-values are indicated above the comparisons.

### Lr-EVs Suppress Intracellular ROS and Stabilize Mitochondrial Gene Expression

Given that SIRT1 closely relationship between cellular redox status and mitochondrial homeostasis, we examined the antioxidant features of Lr-EVs. In cell-free configurations, Lr-EVs displayed powerful, intrinsic radical-scavenging activity against both ABTS radicals (Fig 4A) and DPPH (Fig 4B). *In vitro*, DCFH-DA staining revealed that Lr-EVs robustly quenched TMA-mediated intracellular ROS accumulation, substantially diminishing fluorescence intensity compared to cells exposed to TMA alone (Figs 5 and 6).

**Figure 4.**
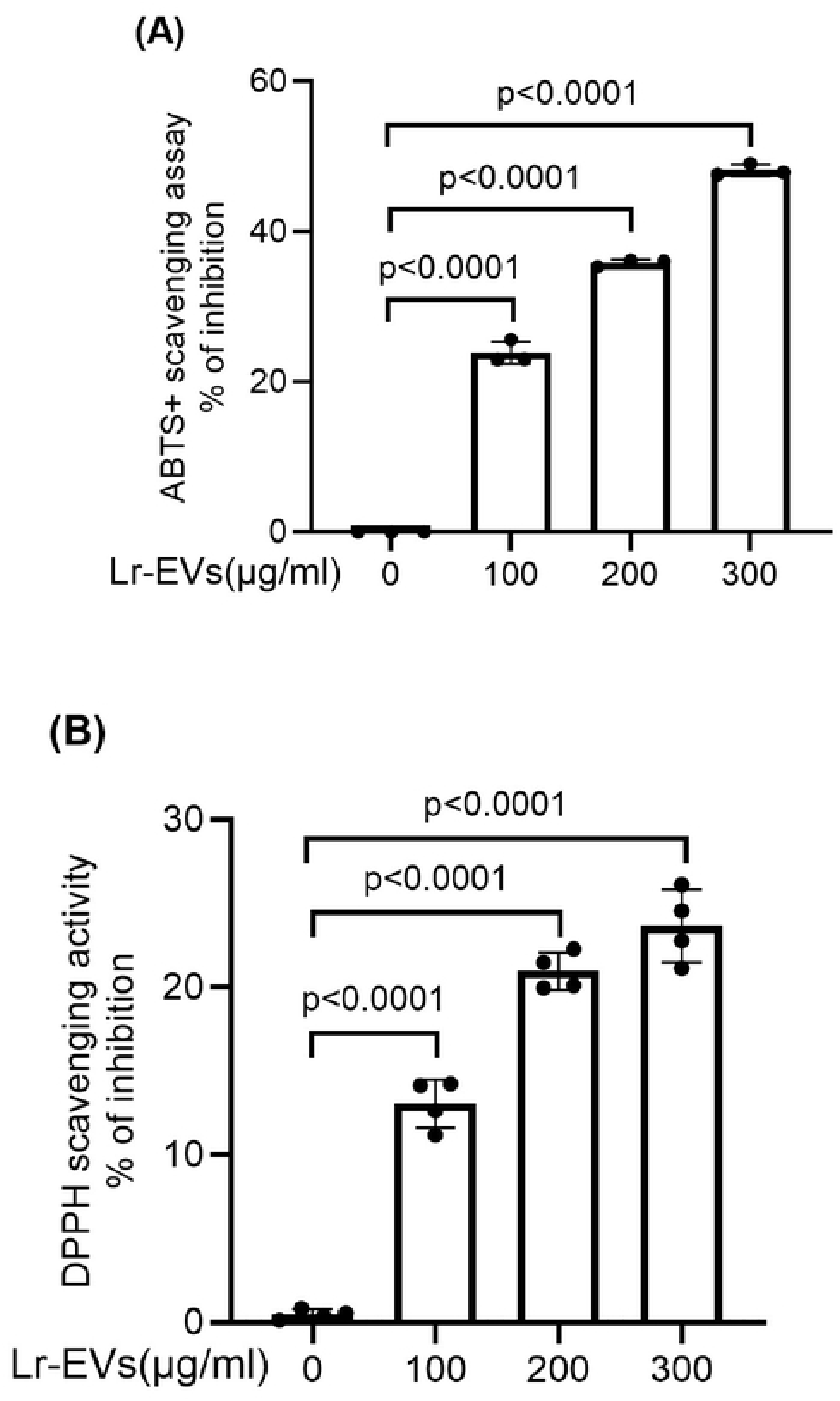
Cell-free radical scavenging capacities of Lr-EVs. Acellular antioxidant performance of varying concentrations of Lr-EVs evaluated by (A) 2,2’-azino-bis(3-ethylbenzothiazoline-6-sulfonic acid (ABTS) radical cation discoloration assay and (B) 2,2-diphenyl-l-picrylhydrazyl (DPPH) radical scavenging assay. *** p < 0.001 vs. untreated control group. Values represent mean ± SD (n = 3).

**Figure 5.**
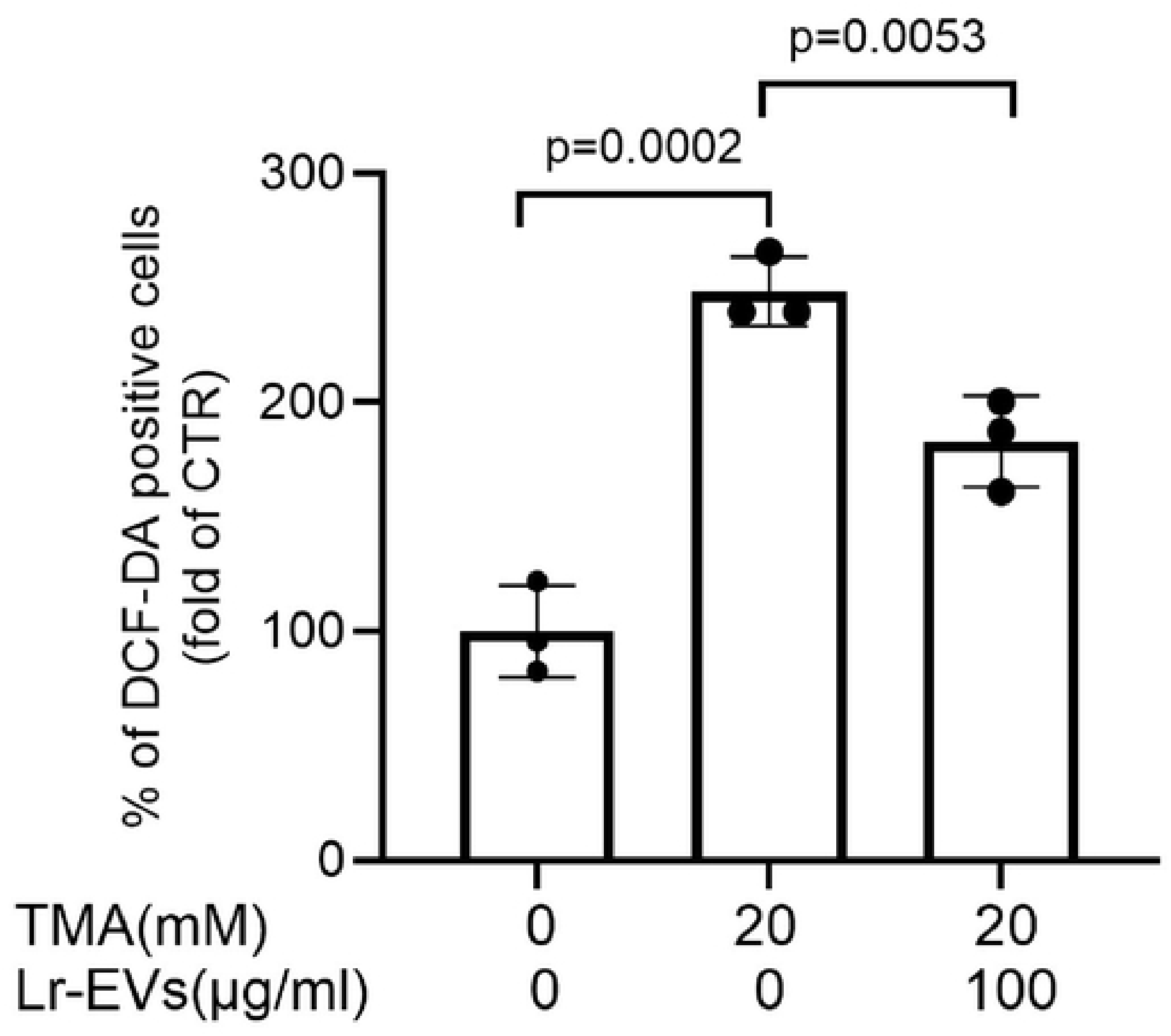
Lr-EVs attenuate TMA-induced intracellular reactive oxygen species **(ROS)** accumulation. Representative fluorescence microscopy images of Caco-2 cells stained with the fluorogenic probe 2,7-Dichlorodihydrofluorescein diacetate (DCF-DA) following 24 h treatments. ** p < 0.01, *** p < 0.001 vs. untreated control group; ## p < 0.01 vs. TMA-treated group. Values represent mean ÷ SD (n = 3).

**Figure 6.**
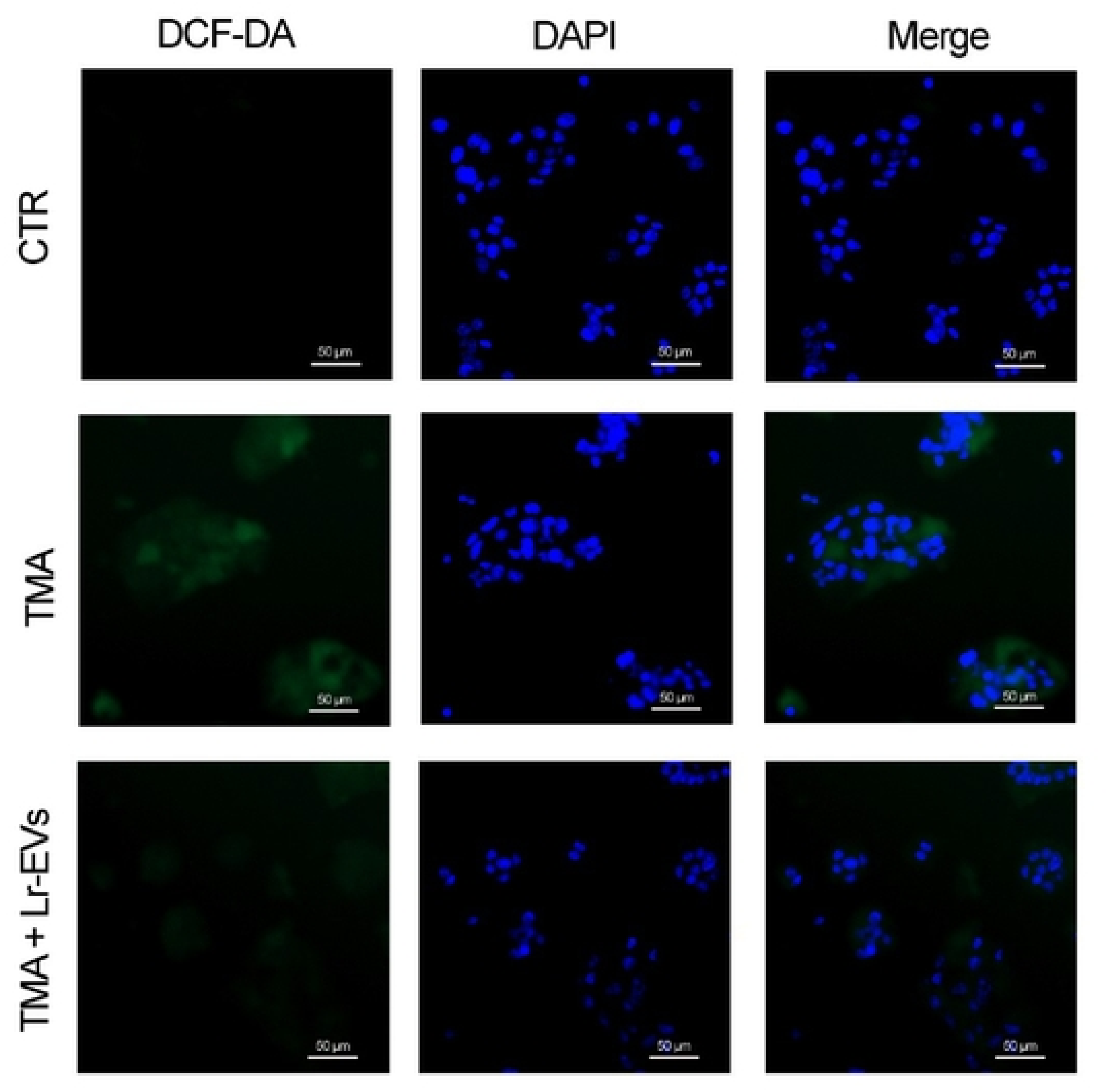
Quantification of intracellular ROS levels in Caco-2 cells. Relative DCF fluorescence intensity quantified from fluorescence images representing intracellular ROS levels across experimental groups. Data are normalized to the control group and shown as mean ± SD (n = 3).

At the transcriptomic level, TMA exposure significantly lowered the expression of essential mitochondrial-encoded electron transport chain (ETC) genes, including ND6, CYTB, CO1, and ATP6. Co-treatment with Lr-EVs successfully stabilized and restored transcript levels for all four genes, demonstrating that Lr-EVs help maintain mitochondrial gene expression under oxidative stress (Fig 7).

**Figure 7.**
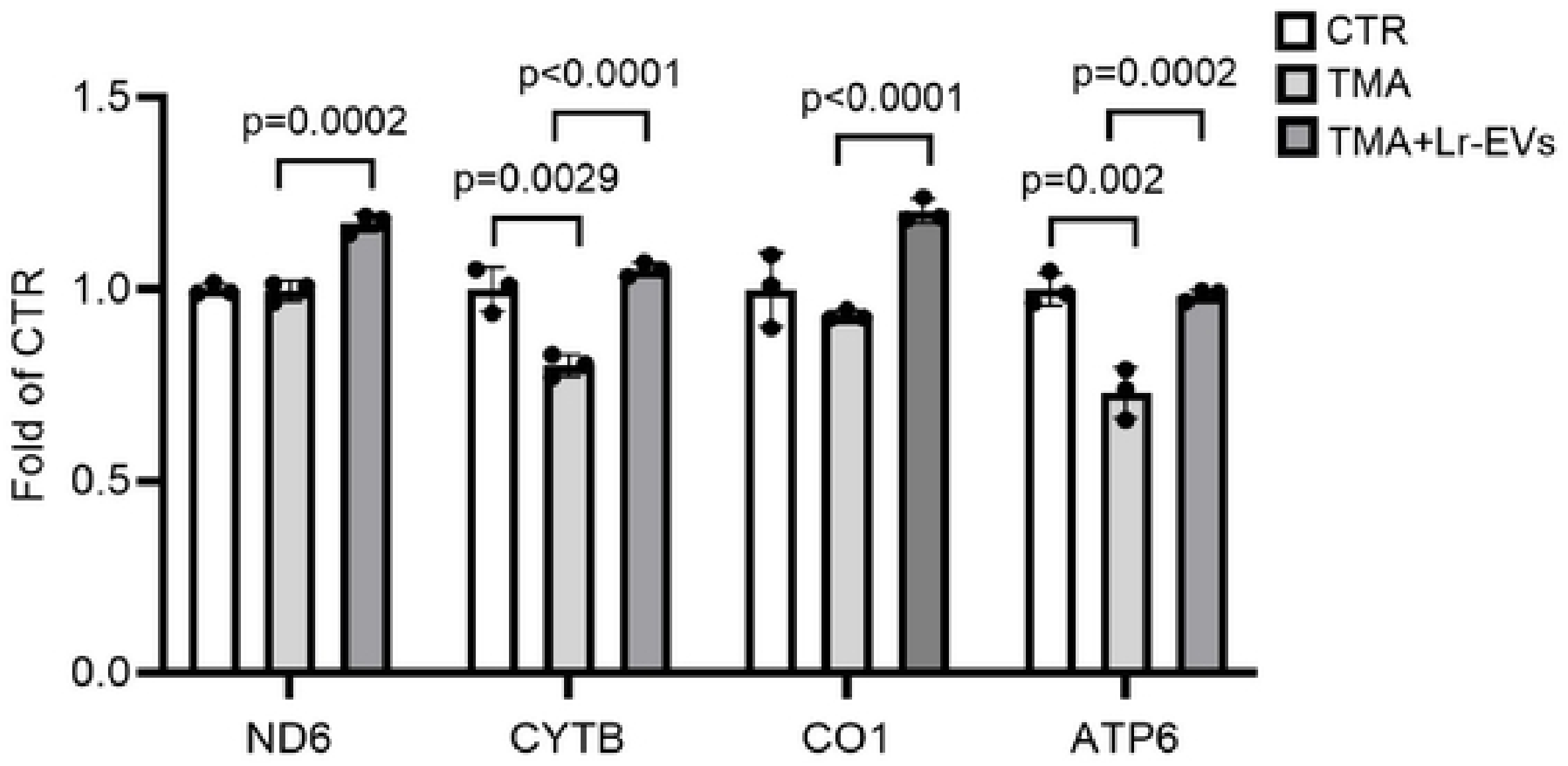
Lr-EVs stabilize mitochondrial respiratory chain gene transcription. Quantitative RT-PCR analysis of mitochondrial-encoded electron transport chain genes in Caco-2 cells following 24 h exposure to TMA and Lr-EVs. Values are normalized to GAPDH and expressed as mean ± SD (n = 3). * p < 0.05, ** p < 0.01, *** p < 0.001 vs. untreated control group; *HUH* p < 0.001 vs. TMA-treated group.

### Lr-EVs Protect the Visual Continuity of Tight Junction ZO-1 Alignment

A damaged intestinal barrier permits the unchecked systemic translocation of luminal microbial metabolites. To evaluate the effects of Lr-EVs on the cellular monolayer organization, the spatial distribution of the tight junction scaffolding protein ZO-1 was monitored. In untreated control monolayers, ZO-1 exhibited a continuous, highly organized, and sharply defined architecture outlining adjacent cell borders. TMA exposure caused extensive breakdown of this network, resulting in a fragmented, discontinuous ZO-1 pattern indicative of severe tight junction disassembly (Fig 8). Remarkably, co-treatment with Lr-EVs fully preserved the spatial architectural continuity and alignment of the tight junction protein ZO-1, maintaining a continuous, well-aligned ZO-1 boundary identical to the control group (Fig 8). This structural reinforcement confirms that Lr-EVs preserve mucosal barrier integrity, potentially blocking localized TMA entry and lowering downstream hepatic TMAO synthesis *in vitro*.

**Figure 8.**
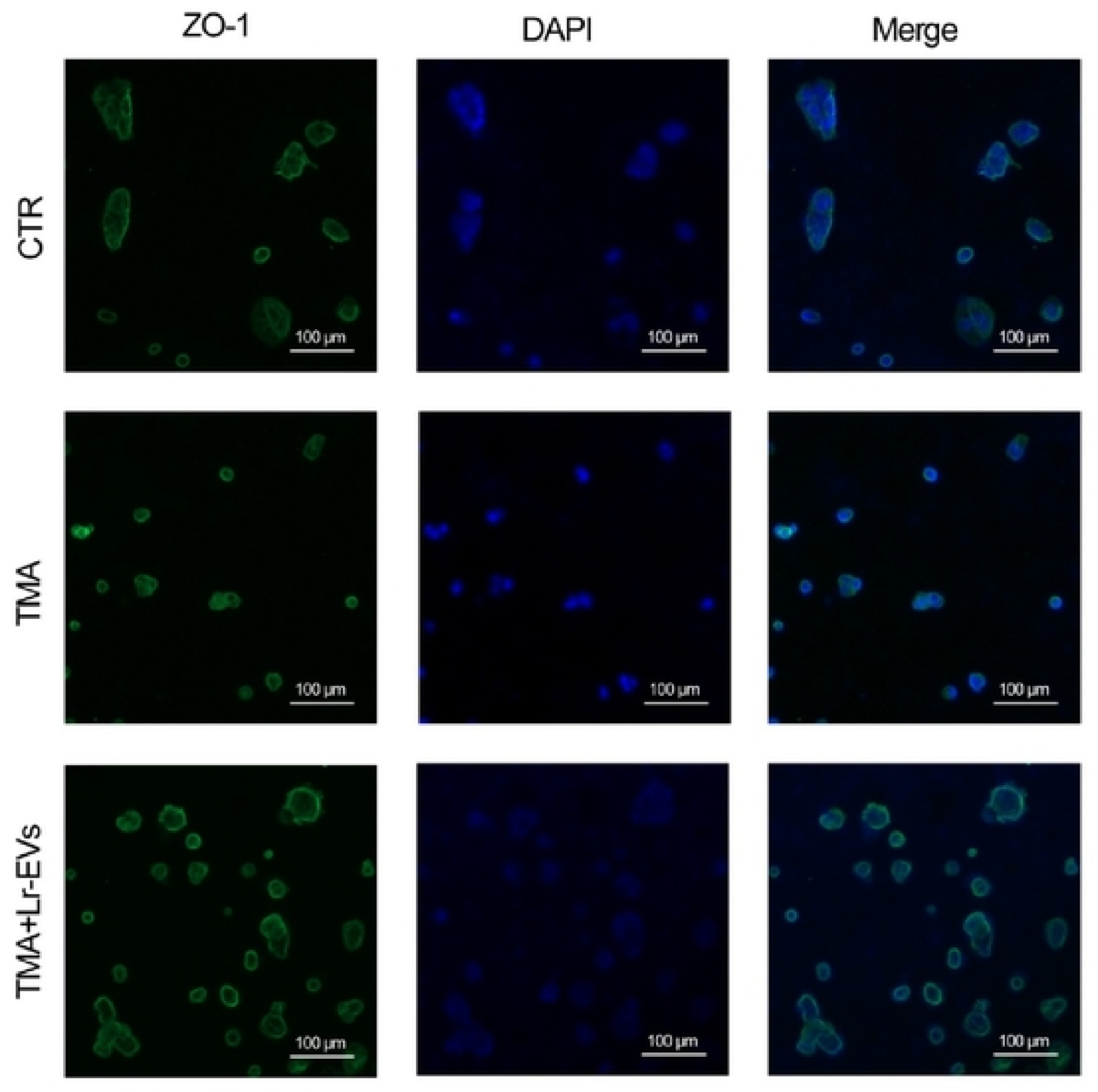
Lr-EVs preserve tight junction ZO-1 structural integrity in Caco-2 monolayers. Representative immunofluorescence staining of tight junction scaffolding protein Zonula Occludens-1 (ZO-1, green fluorescence) in Caco-2 monolayers following 24 h treatment.

## Discussion

This study provides clear evidence that high luminal concentrations of the gut microbiota metabolite TMA induce severe localized cytotoxicity and barrier collapse in intestinal epithelial cells. In aging populations, progressive gut dysbiosis accelerates the conversion of dietary choline and carnitine into TMA (20–22). Our results clarify this pathological process, demonstrating that TMA accumulation disrupts the enterocyte monolayer and compromises tight junction continuity. This structural failure facilitates the paracellular transport of TMA into the mesenteric and portal circulation, driving its subsequent hepatic oxidation into the systemic vascular toxin TMAO. Intercepting this metabolic cascade at the gastrointestinal level therefore represents a powerful approach for the primary prevention of downstream chronic diseases, such as CVD and CKD.

Compared to conventional probiotic interventions, which result in low survival rates due to gastric acid exposure and colonization competition within the host microbiota, extracellular vesicles emerge as a convincing cell-free therapeutic model. We acknowledge that the present study focuses primarily on the rescue effects of Lr-EVs against TMA stress. While isolated Lr-EV-only controls and EV-depleted supernatants were not evaluated in parallel in this baseline screening, these critical controls will be integrated into our next-phase validation to fully dissect the independent bioactivity of the vesicular components. These nanoparticles exhibit superior biostability, low immunogenicity, and the capacity to precisely deliver functional proteins and nucleic acids into recipient cells (23–25). Our findings establish probiotic-derived EVs as an advanced, cell-free alternative to living probiotics. Lr-EVs bypassed the common limitations of live biologics-such as low survival rates in gastric acid and poor niche colonization-and actively stimulated proliferation while protecting against metabolic toxicity. These qualities position probiotic EVs as excellent biochemical delivery vehicles for clinical and nutritional applications.

Sirtuin 1 (SIRT1), an NAD⁺-dependent deacetylase considered a longevity factor, plays an important role in modulating cellular stress responses and metabolic homeostasis. Beyond its metabolic functions, SIRT1 acts as an anti-inflammatory regulator by deacetylating NF-κB, thereby suppressing its transcriptional activity and downregulating downstream pro-inflammatory cytokines (26, 27). Mechanistically, our data reveals that the cytoprotective efficacy of Lr-EVs relies on modulating the SIRT1 axis. SIRT1 functions as a master metabolic switch and anti-inflammatory regulator; it deacetylates the p65 subunit of NF-κB, suppressing its transcriptional activity and reducing downstream pro-inflammatory cytokines like IL-1β. We found that TMA exposure heavily downregulates SIRT1 expression, releasing the brake on inflammation and impairing mitochondrial function. Lr-EV treatment reversed this effect, upregulating SIRT1 and reducing IL-1β levels. It is worth noting that the involvement of the SIRT1 pathway and mitochondrial preservation in this study is characterized as a robust transcriptomic correlation. Although definitive upstream causality awaits further functional verification via western blotting or specific siRNA-mediated silencing, these preliminary qPCR data provide a justified direction for subsequent loss-of-function molecular genetic models. Furthermore, because SIRT1 is an upstream activator of PGC-1α-the central coordinator of mitochondrial biogenesis and respiratory chain integrity (28) this upregulation explains the stabilization of mitochondrial-encoded transcripts (ND6, CYTB, CO1, and ATP6) observed in our study. Thus, Lr-EVs appear to correlated with the transcriptional restoration of the SIRT1 pathway to maintain mitochondrial energetics and suppress downstream cellular damage.

While this study provides clear *in vitro* evidence of Lr-EV-mediated cyto-protection, specific methodological considerations should be highlighted. First, the evaluation of tight junction integrity was performed via the spatial architectural continuity analysis of ZO-1 using high-resolution immunofluorescence microscopy. This approach offers an objective and visually direct assessment of cell-border physical alignment, establishing clear structural preservation even before secondary physiological parameters, such as treneletrical electrical resistance (TEER) or macromolecular flux coefficients (e.g., FITC-dextran), are quantified. Second, the mechanistic link between Lr-EV action and the SIRT1 pathway is currently defined as a robust transcriptomic correlation. Although definitive upstream causality awaits further verification via targeted gene-silencing (siRNA) or pharmacological inhibition assays, this established correlation provides a critical baseline and a justified direction for subsequent loss-of-function genetic blockade models. Lastly, while Caco-2 cell monolayers remain a universally accepted gold standard for preliminary intestinal screening, they lack the multi-lineage cellular complexity of living organisms. Future studies incorporating functional flux measurements, genetic knockdown systems, and complex *in vivo* animal models will be necessary to fully translate the physiological potential of these postbiotic vesicles.

## Conclusion

In conclusion, this *in vitro* study demonstrates that extracellular vesicles isolated from *Lactobacillus rhamnosus* possess intrinsic antioxidant activities and protect human Caco-2 cells against TMA-induced cytotoxicity and oxidative stress. Lr-EVs effectively downregulate the pro-inflammatory transcript IL-1β, stabilize core mitochondrial electron transport chain genes, and protect the structural continuity of the tight junction protein ZO-1, likely via a mechanism involving SIRT1 transcript restoration. These preliminary findings expand our understanding of how probiotic-derived EVs interface with host cells under metabolic challenge and support their further evaluation as cell-free postbiotic candidates in functional food science and healthy aging research.

We acknowledge several limitations in the present study. First, while Lr-EVs demonstrated robust cytoprotective effects against TMA stress, independent Lr-EV-only controls and EV-depleted supernatants were not evaluated in parallel during this initial screening phase. Second, the potential involvement of the SIRT1 signaling axis and mitochondrial preservation is currently established based on transcriptomic (mRNA) correlations. Further loss-of-function investigations using specific siRNA knockdown or Western blotting are warranted to fully elucidate the upstream protein-level causality. Lastly, although Caco-2 cells serve as a well-accepted platform for intestinal cytotoxicity screening, future in vivo animal models will be necessary to translate these preliminary cell-free postbiotic findings into physiological realities.

## Acknowledgements

Financial support for this study was provided by Taipei Medical University - Wan Fang Hospital Joint Research Program under Grant No. 114TMU-WFH-24.

